# Brain-wide continuous functional ultrasound imaging for real-time monitoring of hemodynamics during ischemic stroke

**DOI:** 10.1101/2022.01.19.476904

**Authors:** Clément Brunner, Nielsen Lagumersindez Denis, Karen Gertz, Micheline Grillet, Gabriel Montaldo, Matthias Endres, Alan Urban

**Affiliations:** Neuro-Electronics Research Flanders, Leuven, Belgium; Vlaams Instituut voor Biotechnologie, Leuven, Belgium; Interuniversity microelectronics centre, Leuven, Belgium; Department of neurosciences, KU Leuven, Leuven, Belgium; Department of Neurology and Center for Stroke Research Berlin, Charité-Universitätsmedizin, Berlin, Germany; German Center for Neurodegenerative Diseases (DZNE), Berlin, Germany; German Centre for Cardiovascular Research (DZHK), Berlin, Germany

**Keywords:** Ischemic stroke, functional ultrasound imaging, spreading depolarization, ischemic lesion prediction

## Abstract

Ischemic stroke occurs with no warning, and therefore, very little is known about hemodynamic perturbations in the brain immediately after stroke onset. Here, functional ultrasound imaging was used to monitor variations in relative cerebral blood volume (rCBV) compared to baseline. rCBV levels were analyzed brain-wide and continuously at high spatiotemporal resolution (100μm, 2Hz) until 70mins after stroke onset in rats. We compared two stroke models, with either a permanent occlusion of the middle cerebral artery (MCAo) or a tandem occlusion of both the common carotid and middle cerebral arteries (CCAo+MCAo). We observed a typical hemodynamic pattern, including a quick drop of the rCBV after MCAo, followed by spontaneous reperfusion of several brain regions located in the vicinity of the ischemic core. The severity and location of the ischemia were highly variable between animals. Still, both parameters were, on average, in good agreement with the final ischemic lesion volume measured 24hrs after stroke onset for the MCAo but not the CCAo+MCAo model. For the latter, we observed that the infarct was extended to regions that were initially not ischemic and located rostrally of the ischemic core. These regions strongly colocalize with the origin of transient hemodynamic events associated with spreading depolarizations.

## Introduction

Acute ischemic stroke is most often caused by thrombotic or embolic occlusion of a cerebral artery. It is characterized by a sudden loss of blood circulation to an area of the brain, resulting in a corresponding loss of neurologic function. Ischemic stroke triggers a variable decrease of blood flow in the affected parenchyma that depends on the degree of collateral circulation. In brain regions with poor collaterals, lack of oxygen and glucose may disrupt neuronal activity and ultimately cause a drop in energy metabolism leading to tissue infarction^1,2^. Although both the ischemic core and penumbra are dysfunctional, the penumbra remains viable during a given time window upon restoring blood flow^3^. The penumbra will eventually grow into the ischemic core in the absence of reperfusion. The rate of infarct growth is highly variable between individuals and strongly depends on, e.g., the extensiveness of collateral circulation^4^, the ischemic lesion location, and metabolic factors.

A large number of ischemic stroke models have been developed in rodents to recapitulate the pathophysiology in patients. A particular challenge is the heterogeneity of individual strokes in patients that make it difficult to be modeled in rodents^5^. A significant difference between humans and animal models is the presence of different penumbral and infarct flow thresholds, leading to varying rates and dynamics in infarct evolution^6,7^. The middle cerebral artery (MCA) is commonly affected in ischemic stroke, and hence, occluding this major vessel downstream of the internal carotid artery in rodents may replicate cerebral ischemia in stroke patients. The MCA can be accessed via an approach through either the internal or the external carotid artery and be occluded temporarily or permanently^8,9^. Infarcts induced by this approach often comprise both striatal and cortical damage. Alternatively, blocking the artery distal to the lenticulostriate branches (i.e., distal MCAo) with various strategies (vascular clip^10^, suture^11^, electrocoagulation^12^, photothrombosis^13^, endothelin-1^14^, ferric-chloride^15^, or magnetic nanoparticles^16^) results in a permanent occlusion with mainly cortical infarcts. The pros and cons of stroke models are extensively discussed in the literature by Macrae^17^ and Flury et al.^18^. In short, most models of ischemic stroke involving only the occlusion of the MCA have a large variability^19,20^. Still, it has been shown that a tandem permanent occlusion of the distal MCA and ipsilateral common carotid artery (CCA) can alternatively be used as a surgically simple method for causing large neocortical infarcts with reproducible topography and volume in rat^21^.

Hemodynamics and characteristics of lesion growth following immediately after vessel occlusion are essential issues and represent a new target area for promising therapeutic interventions. Spreading depolarization (SD) describes a slowly propagating wave of intense depolarization that travels across the cortical tissue and is typically coupled with local changes in cerebral blood flow (CBF). SD is associated with a functional hyperemic response when the neurovascular coupling is intact and hypoperfusion when the neurovascular coupling is compromised^22–24^. Moreover, SD may exacerbate ischemic injury^25–29^. The features of SDs induced by ischemia have been widely examined, but studies generally failed to identify their site of elicitation, where they are triggered. Therefore, developing new strategies for monitoring brain perfusion at a large scale and in-depth is crucial to better understand the mechanisms linking perfusion defects and tissue infarction; and improve preclinical drug development^30^. Precise tracking of the hemodynamic responses associated with SDs could help understand their impact on pathophysiology.

The current consensus relies on a multi-modal neuroimaging strategy to provide sufficient information to identify ischemic tissue, biomarkers of changes in tissue metabolism, potentially salvageable tissue, or monitor its reorganization^31,32^. Traditional imaging modalities such as magnetic resonance imaging (MRI), specifically perfusion- and diffusion-weighted imaging (PWI-DWI, pCASL, DSC)^33–35^, positron emission tomography (PET)^32^ and single-photon emission computed tomography (SPECT)^36^ have limited spatial and temporal resolution to resolve vascular responses dynamically. Optical imaging techniques have a high resolution and are well suited to image changes in CBF and blood oxygenation. Laser Doppler flowmetry (LDF)^37,38^ can measure blood flow at a low spatiotemporal resolution. Laser speckle contrast imaging (LSCI)^39^ and optical intrinsic signal imaging (OIS)^40^ are full-field imaging techniques. They can rapidly measure CBF and blood oxygenation, respectively, but only at the surface of the brain. Two-photon microscopy (2PM)^41^ provides vascular morphology and flow information, but it requires exogenous contrast agents. Photoacoustic microscopy (PAM)^42–44^ can measure these parameters but with a limited depth of field. Moreover, the latter two techniques are laser-scanning based, limiting their respective field-of-view and achievable frame-rate, making real-time observation of a larger area challenging.

As an alternative, functional ultrasound imaging used high-frame-rate plane-wave imaging to measure cerebral blood volume (CBV) changes in response to neuronal activity^45–49^. Briefly, functional ultrasound imaging is based on a power Doppler estimator, proportional to the energy of the signals coming from moving scatterers^46^. This technology has a spatial resolution of ~100×300×100μm^3^, a temporal resolution of ~0.1s, and a depth of field of ~1.5cm and is, therefore, ideal for preclinical stroke research^10,45,50,51^. Moreover, it does not require contrast agents and can be adapted to freely moving animals^52–54^.

In this study, we used functional ultrasound imaging to monitor hemodynamic changes during stroke, including perfusion and SDs at the brain-wide scale. We compared two rat stroke models, i.e., a model consisting of permanent occlusion of the distal branch of the middle cerebral artery (MCAo) resulting in small ischemic lesions and a model with a tandem occlusion of the common carotid and middle cerebral arteries (CCAo+MCAo) resulting in larger ischemic lesions typically both comprising large parts of the MCA territory. Hemodynamic changes before were monitored continuously until 70mins after stroke. Data were then registered, segmented, and clustered^55,56^ into 115 anatomical regions, including cortical, thalamic, striatal, and hippocampal areas from a custom rat brain atlas based on Paxinos^57^. Comparing the ischemic territory and the infarcted tissue at 24hrs revealed a mismatch in both its size and location for both stroke models. Notably, we observed that brain regions located rostrally of the ischemic core are not affected by the ischemia up to 70mins after stroke but became part of the ischemic lesion within the 24hrs post-stroke. These regions are also from where SDs emerge quickly after the stroke onset.

## Material and Methods

### Animals

Experimental procedures were approved by the Committee on Animal Care of the Catholic University of Leuven, in accordance with the national guidelines on the use of laboratory animals and the European Union Directive for animal experiments (2010/63/EU). The manuscript was written according to the ARRIVE Essential 10 checklist for reporting animal experiments^58^. Adult male Sprague-Dawley rats (n=24; Janvier Labs, France) with an initial weight between 200-300g were housed in standard ventilated cages and kept in a 12:12hrs reverse dark/light cycle environment at a temperature of 22°C with *ad libitum* access to food and water. To avoid selection bias, animals were randomly distributed in three experimental groups: (i) MCAo, in which the distal branch of the left MCA was permanently occluded (n=10), (ii) CCAo+MCAo, in which the left CCA was permanently occluded shortly before the MCAo (n=10), (iii) a sham group where four rats were imaged as in any other group, but the microvascular clip was inserted next to the MCA without occluding it.

### Cranial window for brain-wide imaging

A cranial window was performed in all rats under isoflurane anesthesia (Iso-Vet, Dechra, Belgium) continuously delivered at 0.6l/min through a nose mask. A mixture of 5% isoflurane in compressed dry air was used to induce anesthesia, subsequently reduced to 2.0-2.5% during surgery and to 1.5% for imaging. Body temperature was monitored using a rectal probe and maintained at 36.5±0.5°C using a digitally controlled heating blanket (CODA, Kent Scientific Corp., USA). Intraperitoneal injection of 5% glucose solution was provided every 2hrs to prevent dehydration. As pre-operative analgesia, Xylocaine (0.5%, AstraZeneca, England) was injected subcutaneously into the head skin. The scalp was shaved and cleaned with iso-betadine before removing the entire dorsal skull. The cranial window extended from bregma (ß) +4.0 to −7.0mm anterior to posterior, laterally 6.0mm to the right side, until the parietal-squamosal suture on the left side to expose the distal branch of the MCA was performed as previously described^10,48,50^. Sterile saline was regularly added during drilling sessions to avoid overheating the tissue. The skull was carefully removed without damaging the dura. Finally, the brain was covered with a low-melting 2% agarose (Sigma-Aldrich, USA) and ultrasound gel (Aquasonic Clear, Parker Laboratories Inc, USA) to ensure a proper acoustic coupling with the ultrasound probe. At the end of the imaging session, Metacam (0.2mg/kg, Boehringer Ingelheim, Canada) was injected subcutaneously for postoperative analgesia, and rats were placed in a warm cage and monitored until waking up.

### MCAo and CCAo+MCAo procedures

Rats from the CCAo+MCAo group underwent CCA dissection before cranial window surgery. Briefly, the rat was placed in a supine position to shave and clean neck hairs. A vertical incision was made on the neck, and the left CCA was carefully dissected from the surrounding tissue and exposed, avoiding damaging the vagus nerve^21,59^. A 3-0 surgical silk thread (Ethicon, France) was placed around the CCA and remained untightened until occlusion during the imaging session. After the imaging section, the neck skin was sutured with a 2-0 surgical silk thread (Ethicon, France). Rats from MCAo and CCAo+MCAo groups were subjected to a permanent MCA occlusion directly during the imaging session using a microvascular clip (#18055-03, micro-serrefine, FST GmbH, Germany) that was manually refined by grinding for optimal serration of the MCAo in adults rats.

### Functional ultrasound imaging

The data acquisition was performed using a functional ultrasound imaging scanner equipped with custom acquisition and processing software described by Brunner et al.^56^. In short, the scanner is composed of a linear ultrasonic transducer (15MHz, 128 elements, Xtech15, Vermon, France) connected to 128-channel emission-reception electronics (Vantage, Verasonics, USA) that are both controlled by a high-performance computing workstation (fUSI-2, AUTC, Estonia). The transducer was motorized (T-LSM200A, Zaber Technologies Inc., Canada) to allow anteroposterior scanning of the brain. Imaging is performed on an anti-vibration table to minimize external sources of vibration. Each coronal Doppler image is 12.8-mm width and 9-mm depth and comprises 300 compound images acquired at 500Hz. Each compound image is computed by adding nine plane-wave (4.5kHz) angles from −12° to 12° with a 3° step. The blood signal was extracted from 300 compound images using a single value decomposition filter and removing the 30 first singular vectors^60^. The Doppler image is computed as the mean intensity of the blood signal in these 300 frames that is an estimator of the cerebral blood volume (CBV)^45,46^. This sequence enables a temporal resolution of 0.6s, an in-plane resolution of 100×110μm, and an off-plane (thickness of the image) of 300μm^56^.

### 2D scan of brain vasculature

Before occlusion, we performed a high-resolution 2D scan of the brain vasculature consisting of 89 coronal planes from bregma (ß) +4.0 to −7.0mm spaced by 125μm. This scan was used for data registration. During the 90-min imaging session, the same brain area was scanned at lower resolution with only 23 planes with a step of 500μm that took approximately 23s to complete. The number of imaging planes was chosen to maximize the number of brain volumes scanned per min while preserving a good resolution in the anteroposterior axis. For the MCAo group, a 20-min baseline was recorded before occluding the MCA, whereas the CCAo+MCAo group has a 10-min baseline before CCAo, followed by the MCAo 10mins later. Notably, the procedures were performed live with a real-time display of the Doppler images, allowing for monitoring the brain perfusion and the direct confirmation of the CCA and/or MCA occlusions (Movies 1 to 3). The imaging session was stopped after 90mins of recording.

### Registration and segmentation

We developed a custom digital rat brain atlas for registration of the data based on the one stereotaxic atlas of Paxinos^57^. In short, the spatial transformation matrix was computed on the high-resolution 2D scan that was manually aligned on the atlas by affine transformation (i.e., translation, rotation, and scaling). This matrix was then applied to the low-resolution 2D scan. The dataset was segmented into 115 anatomical regions/hemispheres that were subsampled from the 332 brain regions of the original atlas (see Supplementary Table 1). The hemodynamic signals were averaged in each area. The software for data registration and segmentation is open-access here^61^. The unrolled-cortex representation corresponds to the maximum intensity projection of the signal located 250μm under the cortical surface (see Supplementary Figure 1).

### Relative cerebral blood volume (rCBV)

We used the relative cerebral blood volume (rCBV; expressed in % compared to baseline) to analyze ischemia defined as the signal in each voxel compared to its average level during the baseline period (Figure 1). After registration and segmentation, the rCBV signal was averaged in each individual region.

**Figure 1.**
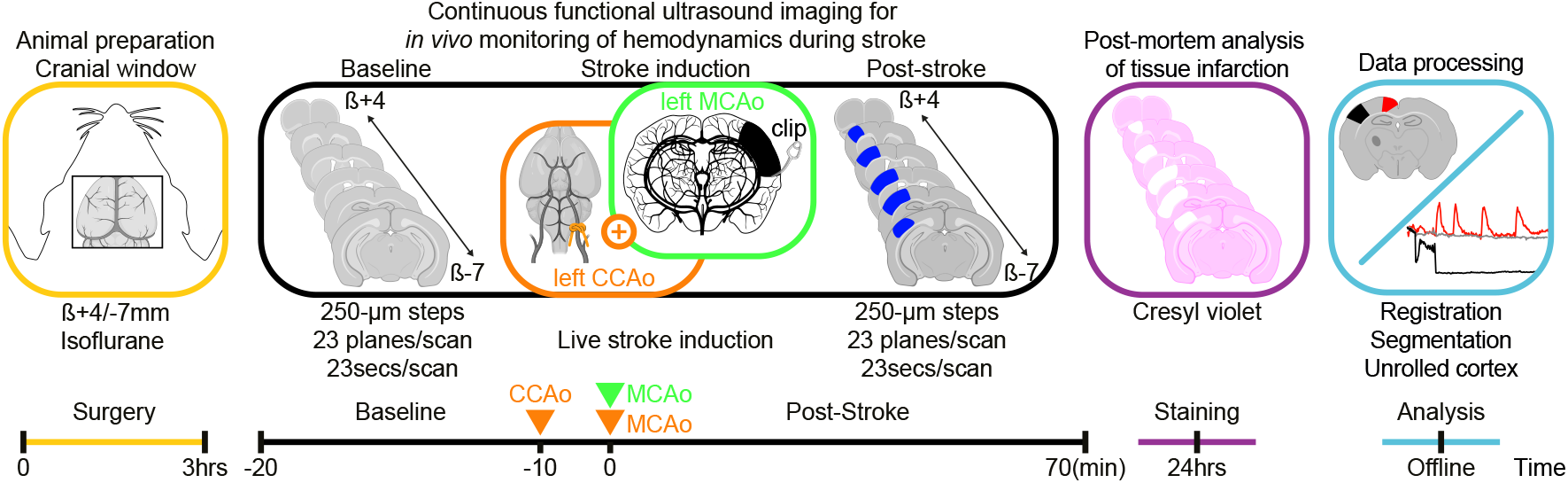
Experimental workflow and timeline. From left to right. An 11×13-mm cranial window was performed to access the whole brain by functional ultrasound imaging before occluding the middle cerebral artery occlusion (MCAo) or MCAo combined with the common carotid artery (CCAo+MCAo). Image acquisition has been performed continuously for 70mins by scanning the brain repeatedly in the anteroposterior direction (23secs/scan) before, during, and after stroke onset. After the experiments, rats returned to their home cage and were euthanized 24hrs after occlusion to quantify the infarct size using cresyl violet staining. We developed a digital version of the rat Paxinos atlas^57^ for the registration, segmentation, and temporal analysis of ischemia using a dedicated software solution^61^.

### Clustering of the rCBV loss

Once registered and segmented, the brain regions were clustered based on the average rCBV loss after the MCA occlusion and sorted from the most significant loss of rCBV signal to the smallest (from 1 to 115) as compared to baseline level. The list of the brain regions clustered can be found in Supplementary Table 1.

### Analysis of spreading depolarizations

The detection of spreading depolarizations (SDs) was performed based on the temporal analysis of the rCBV signal. The rCBV signal was averaged in an area of interest (10×10 voxels) located in the left side cortex, laterally to the ischemic territory. An SD is defined as a transient increase of rCBV signal above 50% compared to baseline. This procedure allowed to measure the occurrence of SDs over the recording period and their propagation pattern in the field of view. The velocity of each SDs was calculated as the time spent to propagate between two areas of interest located in the left side cortex, along with the ischemic territory. The origin of each SDs was located by tracking back the trajectory to the region of the first detection of the transient increase of rCBV. The occurrence, the propagation pattern, and the velocity of SDs could be efficiently visualized using the unrolled-cortex projection as presented in Movie 3.

### Histopathology

Rats were killed 24hrs after the occlusion for histological analysis of the infarcted tissue. Rats received a lethal injection of pentobarbital (100mg/kg i.p. Dolethal, Vetoquinol, France). Using a peristaltic pump, they were transcardially perfused with phosphate-buffered saline followed by 4% paraformaldehyde (Sigma-Aldrich, USA). Brains were collected and post-fixed overnight. 50-μm thick coronal brain sections across the MCA territory were sliced on a vibratome (VT1000S, Leica Microsystems, Germany) and analyzed using the cresyl violet (Electron Microscopy Sciences, USA) staining procedure. Slices were mounted with DPX mounting medium (Sigma-Aldrich, USA) and scanned using a brightfield microscope. Ischemic lesion areas were manually delineated and measured by two researchers independently in double-blind conditions using the Fiji software^62^. Brain slices were then registered on the rat atlas utilizing a set of landmarks, including the overall size of the slice, the cortical thickness, the positions of ventricles, and the position/size of the hippocampus. The ischemic lesion was projected in the unroll-cortex using the same approach as defined previously (see above and Supplementary Figure 1).

### Statistical analysis

The dataset successfully passed Shapiro-Wilk and Kolmogorov-Smirnov tests (significance level α=0.05) for normal distribution before being subjected to statistical analysis, see details in results and figure captions. All data are shown as mean±standart deviation(sd). Statistical analysis were performed using Prism9.3.1 (GraphPad Software).

## Results

### Animals

Three rats from the MCAo group were excluded from the analysis because we observed an extensive *postmortem* hemorrhage extending in the cortex and subcortical regions. It was probably caused by a displacement of the microvascular clip during the 24hrs recovery period. Therefore, the number of rats in this group was reduced to n=7 for the analysis.

### Brain-wide continuous monitoring of hemodynamics

The functional ultrasound imaging scanner allows repeated 2D scans in the anteroposterior direction. For the first time, we demonstrate the applicability of this technology for large-scale and continuous monitoring of hemodynamics in deep brain tissue before, during, and after stroke onset. After data acquisition, every 23 cross-sections spaced of 500μm were registered and segmented on a custom-developed digital rat atlas (see Materials and Methods) to provide a volumetric and dynamic view of the changes in perfusion at high spatiotemporal resolution.

Typical plots of the temporal evolution of the hemodynamic signal are shown in Figure 2(a) for both the MCAo (green plot) and CCAo+MCAo (orange plot) stroke models. In this example, the variations in the cerebral perfusion were measured as rCBV compared to the baseline level in two regions of interest located either in the ischemic core in the left hemisphere (S1BF) or in the opposite control hemisphere (black ROI and plots). As shown, for the MCAo stroke model (Figure 2(a), green plot), the rCBV signal dropped immediately by ~ −50% after MCAo and remained around this value during the entire duration of the experiment. On the contrary, we observed a two-step and a more profound decrease of the rCBV signal (Figure 2(a), orange line) in the CCAo+MCAo stroke model. First, we observed a transient drop of the rCBV to approximately −60% of the baseline level caused by the CCA occlusion that quickly recovered but not up to the initial values. Then, the MCAo triggers a massive reduction of ~ −95% of the rCBV that remains at this level during the entire recording. Interestingly, in the CCAo+MCAo stroke model, the contralesional hemisphere also shows a transient drop of ~ −15-20% of the rCBV signal after CCAo that is not exacerbated after MCAo (Figure 2(a), black line). Note that the rCBV in the control ROI located in the opposite hemisphere remains stable all along with the experiment in both stroke models (Figure 2(a), dark line).

**Figure 2.**
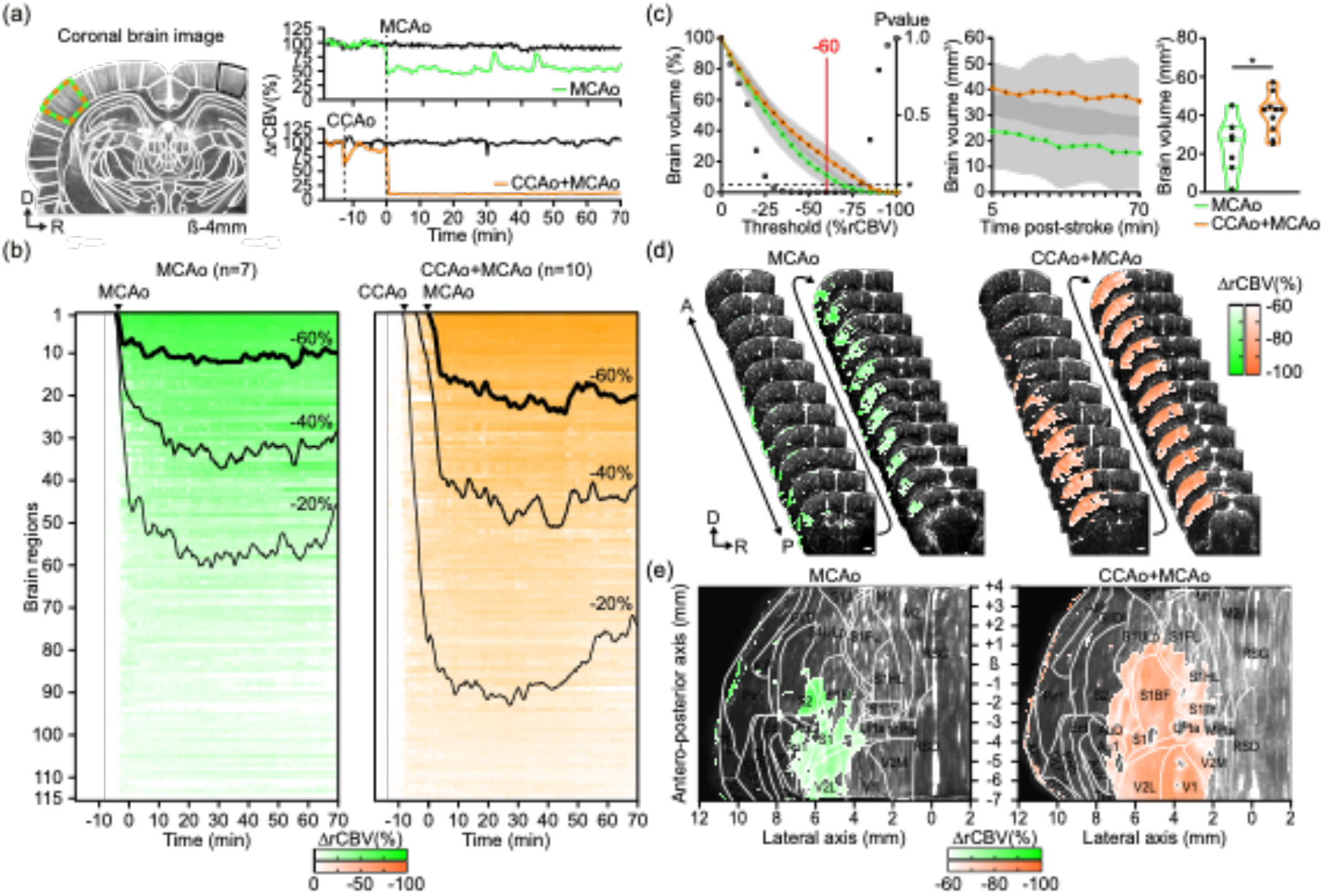
Brain-wide continuous and real-time monitoring of hemodynamics during ischemic stroke. (a) A typical image of the brain microvasculature for one coronal cross-section from the 2D scan before the stroke. Only the left hemisphere is imaged entirely here because of the too-small size of the ultrasound transducer used for this study. All images were registered and segmented based on a digital version of the rat Paxinos atlas^57^ (white outlines). Time series plots of the average signal in the primary somatosensory barrel-field cortex (S1BF; green and orange region) in the affected hemisphere and the control ROI located in the opposite hemisphere (black region) for each stroke model (MCAo in green; CCAo+MCAo in orange). (b) Global hemodynamic changes (rCBV) in 115 regions located in the ischemic hemisphere for both the MCAo (left and green panel) and CCAo+MCAo groups (right and orange panel). Regions were clustered according to the extent of rCBV signal loss (from more to less pronounced) (see Supplementary Table 1). (c) Left to right. Plot showing the total volume of brain region corresponding to a given rCBV decrease (%; mean±sd, multiple unpaired t-test, *p<0.05; left panel). Evolution of brain volume with 60% rCBV reduction during the 70-min post-stroke period (mean±sd; middle panel). Brain volume with a rCBV reduction of −60% at 5mins after stroke onset at a signal threshold of 60% (Unpaired t-test, *p=0.013; right panel) between MCAo (green, n=7) and CCAo+MCAo groups (orange, n=10). (d) Typical 2D brain scan of hemodynamics showing the loss in rCBV induced by either MCAo (left panel) or CCAo+MCAo (right panel). 22 cross-sections out of 23 are represented here. (e) Unrolled-cortex projection (see Supplementary Figure 1) showing the loss in rCBV induced by either MCAo (left panel) or CCAo+MCAo (right panel). D: dorsal, R: right, A: anterior, P: posterior, ß: Bregma reference point. Scale bars: 100μm.

Figure 2(b) provides a detailed view of hemodynamic changes in 115 brain regions in the affected hemisphere (See Supplementary Table 1) averaged for all animals in each group (MCAo group, n=7; CCAo+MCAo group, n=10). Each region is sorted from the most significant loss of rCBV signal to the smallest (from 1 to 115).

Three rCBV levels (−60, −40, and −20%) were defined to better compare the MCAo and CCAo+MCAo stroke models. At the level above −60% rCBV, we noticed that more regions were affected in the CCAo+MCAo than in the MCAo stroke model (~20 vs. ~10, respectively), confirming the cumulative effect of MCAo+CCAo on the reduction of the perfusion. Note that all these regions were exclusively located in the cortex. Between −40 and −60%, approximately the same number of regions (~30) were affected in the two models, but we observed that those regions tend to be slightly reperfused naturally over time (Figure 2(b)). Furthermore, we identified 50-60 regions located in the cortex with a drop of rCBV signal below 20% of the baseline level in the MCAo group, whereas 80-90 regions were affected for the same levels in the MCAo+CCAo group also including subcortical structures (Figure 2(b) and Supplementary Table 1).

To evaluate the relevance of the rCBV signal to precisely assess the perfusion status and differentiate each stroke model, we quantified the affected brain volume (in mm^3^) for various levels of rCBV decrease (Figure 2(c)). We observed that ischemic volume differs significantly between the MCAo and the CCAo+MCAo groups for values ranging from −30 to −75% (Figure 2(c)), left panel; mean±sd, Unpaired t-test, *p<0.05). The most significant difference between the two groups was observed for the value −60% that was chosen for further analysis (Figure 2(c) to (e)).

The total volume of brain tissue showing an rCBV reduction of −60% was then analyzed during the entire imaging session. We observed that the volume of ischemic tissue for the CCAo+MCAo group was 41±10mm^3^ and did not show a significant change between the MCA occlusion and the end of the imaging period (35±14mm^3^; Figure 2(c), middle panel; mean±sd; p>0.44, GLM-based repeated measures one-way ANOVA). On the contrary, the volume of tissue affected for the MCAo group tends to shrink over time from 24±14mm^3^ quickly after MCAo to 15±14mm^3^ after 70mins afterward (Figure 2(c), middle panel; mean±sd; p<0.0001, GLM-based repeated measures one-way ANOVA). Interestingly, the differences between the two groups for an rCBV level of −60% were already statistically significantly different at 5mins after stroke onset. It became even more pronounced during the 70mins imaging period (Figure 2(c), right panel; p=0.013; Unpaired t-test).

Figure 2(d) presents a typical case for each model in which we overlaid the ischemic territory on top of each microvascular image collected during brain scans for each stroke model. We projected and unrolled the cortical regions into a flat surface overlaid with the anatomical landmarks from the reference rat brain atlas to appreciate the differences better. Such a view is advantageous for accurately visualizing the regions with reduced rCBV, their spatial extent (Figure 2(e) and Supplementary Figure 1), and comparing animals and models.

### A large part of the infarct is located in regions not ischemic at 70mins after stroke onset

Cresyl violet staining of coronal brain slices was performed 24hrs after the experiments. A typical example is presented in Figure 3(a). It shows that the infarcted area is localized only in the cortex in both stroke models, but MCAo caused a much smaller and less extensive lesion (Figure 3(a), left panel; green area) than CCAo+MCAo (Figure 3(a), right panel; orange area).

**Figure 3.**
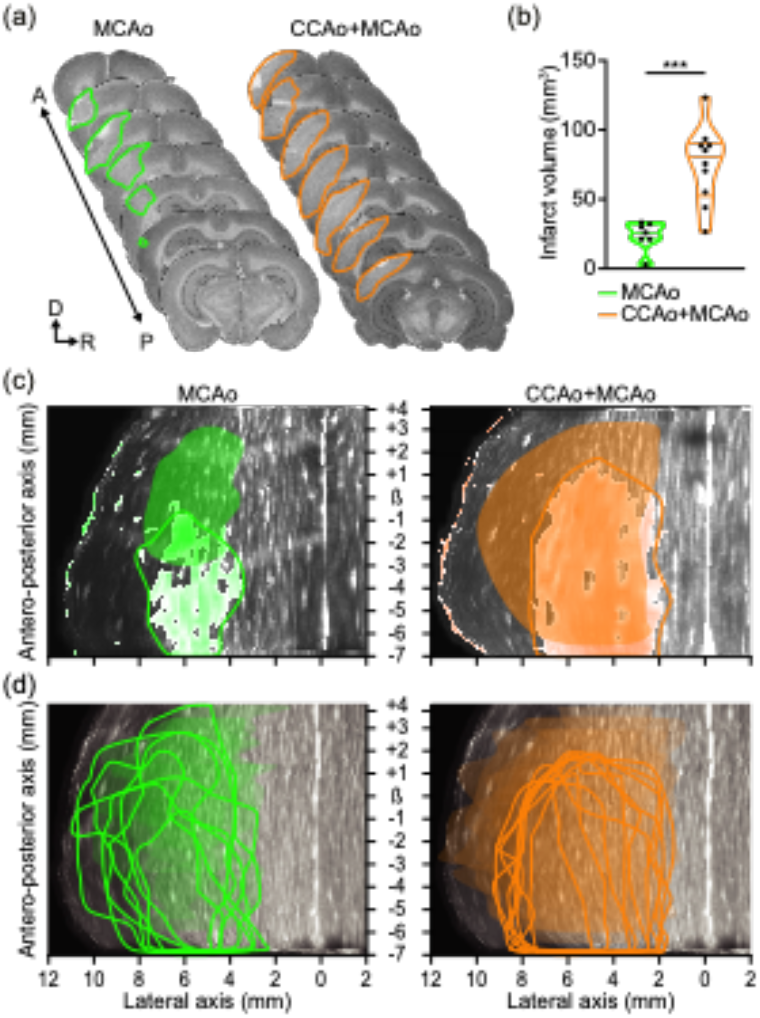
Size and location of the region with a low rCBV signal at 70mins do partially overlap with the infarct at 24hrs. (a) Typical rat brain cross-section stained by cresyl violet to evaluate the infarct size at 24hrs after MCAo (left) or CCAo+MCAo (right). The infarcted territory is highlighted for each stroke model (green and orange, respectively). (b) Comparison of the infarct volume (mm^3^) between the two models, showing the CCAo+MCAo (orange) display a statistically more extensive infarct than the MCAo group (green; Unpaired t-test, ***p=0.0003). (c) The infarct (colored halos) and the region with low rCBV levels (colored circles) are overlaid on the unrolled-cortex projection for one typical rat. (d) Same as in (c) but for all rats used in the study. D: dorsal, R: right, A: anterior, P: posterior, ß: Bregma reference point.

The statistical analysis of all rats confirmed that the infarct volume is significantly larger for the CCAo+MCAo group (75±28mm^3^, n=10, mean±sd; Figure 3(b), in orange) when compared to the MCAo group (24±10mm^3^, n=7, mean±sd; Figure 3(b), in green; ***p=0.0003, unpaired t-test). Note that these results are in agreement with rCBV data, also showing a larger ischemic territory in the CCAo+MCAo group. Taken together, this demonstrates that functional ultrasound images acquired at early time points after stroke onset (from 5 to 70mins) have an excellent predictive value to predict brain infarction. To confirm this hypothesis, we compared the results obtained by histopathology with those from functional ultrasound imaging in each stroke model that were both presented using the same unrolled-cortex projection as described previously (see Supplementary Figure 1). Figure 3(c) shows a typical case for each group. We observed in the MCAo model (Figure 3(c), left panel) that the infarcted region (green circle) only partially coincides with regions showing reduced rCBV (green halo). Interestingly, much of the infarcted tissue is located rostrally to the region with low rCBV at 70mins, in an area that was not initially ischemic. On the contrary, we observed a considerable overlap between the infarct region and the initially ischemic region at 70mins in the CCAo+MCAo model (Figure 3(c), right panel) even though, here again, non-overlapping regions were observed in the anterior part of the brain. Nevertheless, when total infarct size was compared with the initial ischemic territory, it was preserved in the MCAo but not in the CCAo+MCAo model, in which the infarct volume was much smaller than the initial ischemic region.

Although we observed large variability in terms of the location and size of the infarct at 24hrs after stroke compared with the ischemic areas at 70mins after stroke onset for each animal (Figure 3), these results were confirmed at the group level, showing that the infarcted territory extends rostrally to the ischemic region by 1.53±1.68 and 1.37±0.66mm, respectively in MCAo and CCAo+MCAo groups (p=0.49, Unpaired t-test). This mismatch is consistent for the two groups (6/7 for MCAo and 7/7 for CCAo+MCAo rats). Additionally, we noticed that the MCAo group also shows, on average, a mismatch between the position of the infarct and the areas of low rCBV in the posterior part of the brain, where we previously observed good reperfusion during the 70-min period of the functional ultrasound imaging recording (Figure 2(b) and (c)). In both groups, these results indicate that the lack of blood supply may not be the only factor contributing to the infarction of the brain.

These findings prompted us to evaluate other important factors, such as SDs that have been shown to exacerbate focal ischemic injury by converting zones of the viable but non-functional ischemic penumbra to the core region beyond rescue^63^.

### Spreading depolarizations start quickly after MCAo

As shown in previous studies, functional ultrasound imaging is suitable for whole-brain tracking of transient hyperemic events associated with SDs^45,64^. The imaging protocol that we developed for this study allows for detection, monitoring, and quantification of SDs in real-time before, during, and after stroke onset. The spatiotemporal profile of SDs was extracted by averaging the rCBV signal in all voxels from the retrosplenial granular cortex (RSG) that is located in the vicinity of the region with the low level of rCBV (Figure 4(a), left panel). As already observed in previous studies, SDs were triggering transient hyperemia characterized by a rapid and massive increase of the rCBV signal (>+150% rCBV), followed by a sustained reduction of the baseline level (Figure 4(a), right panel, and Movies 2 and 3). We did not observe a significant difference in the number (4.3±2.3 and 4.7±2.9, mean±sd, p=0.7557, Unpaired t-test) and the frequency (1 SD each 17.5mins and 15.2mins, central frequency) of SDs between the MCAo and CCAo+MCAo groups (Figure 4(b)). A detailed analysis of one individual animal revealed that all SDs are generated within the same brain region (Movie 3). The sham controls had a maximum of 1 SD occurring in the first 5mins but did not show any infarcted tissue in the *postmortem* analysis (n=4, data not shown). It has been reported that the initial SD could be due to the mechanical pressure during the mechanical clipping to occlude the MCA as previously reported^65^. If we exclude this initial SD from our analysis, we observed that the second SD appears rapidly within ~10mins for both models (MCAo: 6±11mins, CCAo+MCAo: 11±9mins). The group analysis of functional ultrasound imaging data for all animals revealed that most SDs were originating from the primary somatosensory cortex (72.2% for MCAo versus 83.3% for CCAo+MCAo) between S1ULp and S1FL (centroid coordinates: ß+1.5/2mm, Lateral 4.5/5mm) in the close vicinity of the area with low rCBV signal (Figure 4(c)). SDs are propagating at a constant velocity. We did not observe differences in the velocity of SDs between MCAo and CCAo+MCAo groups (respectively 5.8±1.0 and 6.4±2.4mm/min, mean±sd; p=0.5683, Unpaired t-test; Figure 4(d)), but the values are in agreement with others studies^66^. Figure 4(e) presents a typical color-coded map overlaid on the unroll cortex projection showing the trajectory and progression of a standard SD from the anterior to the posterior part of the brain by contouring the ischemic territory (Figure 4(e) and Movie 3).

**Figure 4.**
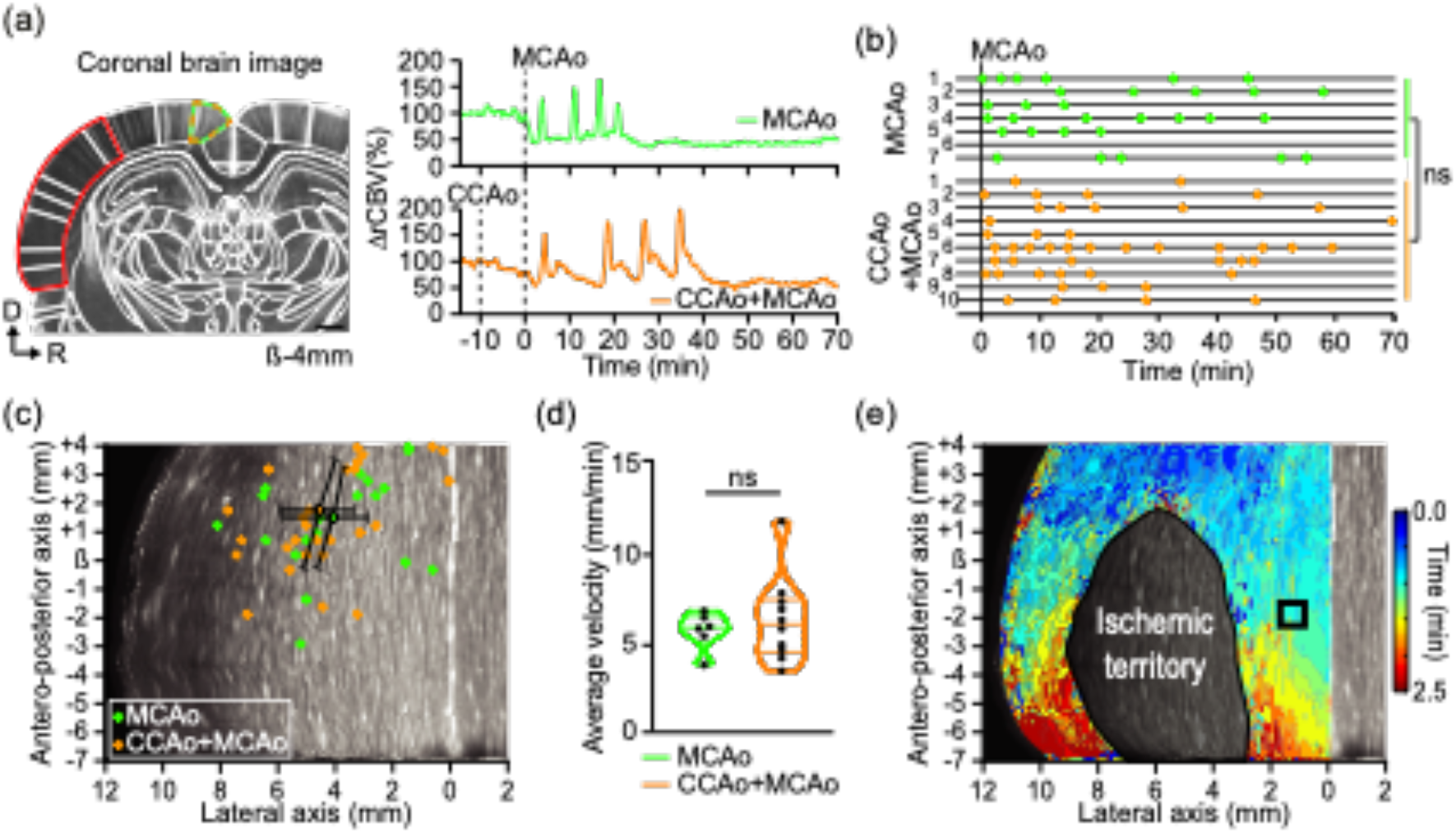
Real-time monitoring of spreading depolarizations using functional ultrasound imaging. (a) A coronal crosssection extracted from the 2D functional ultrasound imaging scan after stroke onset shows the substantial reduction of the rCBV signal in the cortex (red area). The time series plot of the average hemodynamic signal (rCBV) in the retrosplenial granular cortex (green and orange dotted line) from the ischemic hemisphere for each stroke model (MCAo in green; CCAo+MCAo in orange). (b) Monitoring of SDs. Each horizontal line represents one animal; each dot corresponds to the transient hemodynamic increase associated with an SD. The average SDs are not statistically different between the two groups (Unpaired t-test, ns=0.7557). (c) Location of the epicenter of SDs for the MCAo (green dots) and CCAo+MCAo (orange dots) models with respective centroids (mean±sd). (d) A plot of the distribution of the average velocity of SDs for MCAo (green) and CCAo+MCAo models (orange). Each point represents one rat. The average SD velocity is not statistically different between the two groups (Unpaired t-test, ns=0.5683). (e) Color-coded map showing the propagation of an SD in a concentric manner around the ischemic core from its epicenter located in the anterior part of the brain to the posterior part. D: dorsal, R: right, ß: Bregma reference point. Scale bar: 100μm.

## Discussion

To recapitulate the diversity of stroke observed in patients, various experimental models and surgical procedures are employed in rodents, including transient or permanent occlusion of MCA and/or CCA with a filament, photo- or chemo-thrombosis^15–18^. However, each strategy is often associated with complex and multifaceted stroke mechanisms, limiting their comparability^17,18^.

In this work, we used two stroke models, in nearly identical conditions, using permanent occlusion (with a vascular micro-clip) of the MCAo combined with or without CCAo (ligature) to reduce collateral reperfusion, causing either small or large tissue infarction, respectively. This procedure was chosen to allow a fair comparison of brain-wide hemodynamic monitoring with functional ultrasound imaging between the two models, from early time point after stroke onset and during 70mins afterward.

Several imaging technologies such as magnetic resonance angiography (MRA), computed tomography perfusion, contrast-enhanced CT angiography, and transcranial Doppler sonography (TCD) are used to seek evidence of plausible stroke mechanisms and assess the efficiency of recanalization strategies^67^.

However, hemodynamic patterns are highly variable since either improvements or deterioration related to dynamic changes in brain perfusion may occur early during the clinical course of ischemic stroke. These changes are often associated with spontaneous thrombolysis, re-occlusion, micro-embolism, thrombus propagation, and collateralization^68^. CT perfusion can be performed within 30mins after hospital admission, but most often information on brain hemodynamics is available only after several hours to days after the stroke onset, which may be too late for significantly improving the diagnosis, care, and prognosis of stroke patients^69^. Early, continuous and repeatable monitoring of cerebral hemodynamics may offer new insights into acute ischemic stroke pathogenesis. In theory, precise recording of brain perfusion may even guide therapeutic interventions to prevent neurological deterioration or plan conservative treatment in response to particular hemodynamic patterns.

This study demonstrated that functional ultrasound imaging is a suitable technology for continuous monitoring of brain-wide hemodynamics during stroke, allowing for real-time assessment of the ischemic insult^10,50,51^. For the first time, we analyzed the changes in rCBV at high spatiotemporal resolution in both MCAo and CCAo+MCAo stroke rat models, which are used mainly for preclinical stroke research^70^. Technically, we performed repeated 2D scans in the anteroposterior direction (23 positions, 11×12.8×9-mm^3^ total brain volume) to continuously monitor the rCBV signal from stroke onset to 70mins afterward. Functional ultrasound imaging data were then registered and segmented across 115 regions based on a digital version of the rat Paxinos atlas^57^ using a custom software solution previously developed for mice^55,56,71^. This analytics pipeline is open-access and available for research purposes^61^. It is a powerful tool for efficiently comparing hemodynamic changes in the entire brain between animals of the same group and between groups. Volumetric (Figure 2(d)) and cortical projection (Figure 2(e)) maps of the ischemic territory for each rat were reconstructed from the stack of the 2D slices to analyze the variability between animals and to compare it with histopathological data on the infarct (Figure 3(d)).

As expected, we observed that the rCBV loss and its spatial extend are more pronounced in the CCAo+MCAo than in the MCAo model, probably due to a reduction of the collateral flow in MCA territory after CCAo^72–74^. We defined four thresholds (above −60%, −40%, −20%, and below as compared to the baseline) to compare the two models and showed that the identity and number of regions in each category strongly differ between the two models. Moreover, we observed complex hemodynamic patterns throughout the entire imaging period, including pre-occlusion, early before and after CCAo+MCAo, to 70mins afterward. Thanks to the high spatiotemporal resolution of functional ultrasound imaging and its ability to image hemodynamics in deep brain regions, we measured decreasing rCBV levels in the ischemic territory from the cortical surface to deep subcortical regions with typical rCBV values ranging from −100 to 60% in the cortex, ~ −40% in the hippocampus and ~ −20% in the striatum (Figure 2(b)).

Even if we observed considerable variability in the size and location of the ischemic regions between animals (Figure 3(d)), we demonstrated that the MCAo and CCAo+MCAo stroke models are statistically significantly different when comparing the number of regions with an rCBV level below −60% (Figure 2(c) to (e)). For this threshold, we observed that all ischemic regions were located in the cortex (Figure 2(b)) and that the volume of tissue with an rCBV value below −60% progressively decreases from 24mm^3^ 5mins after stroke onset to 15mm^3^, 70mins post-stroke in the MCAo model (pvalue=0.0117, Paired t-test; Figure 2(c); middle panel). Such an effect was not observed for the CCAo+MCAo model (pvalue=0.1022, Paired t-test; Figure 2(c), middle panel). When comparing only the regions with an rCBV level above −60%, we showed a slow but continuous perfusion increase following the MCAo in both stroke models (Figure 2(b)).

We quantified the infarct size and found it statistically significantly smaller for the MCAo than for the CCAo+MCAo model (25 vs. 75mm^3^) despite considerable inter animal variability in both models, as previously reported for various preclinical stroke models^75^ and patients^76^. Notably, we observed a significant dispersion of infarct sizes in the CCAo+MCAo than in the MCAo model, possibly because of spontaneous reperfusion that may occur outside the imaging period (Figure 2(b)), even though this model has been shown to provide more reproducible stroke lesions^21,77,78^. Notably, the infarct volume measured at 24hrs *postmortem* did match those of low rCBV measured *in vivo* using functional ultrasound imaging for the MCAo but not the CCAo+MCAo model. For the latter, the infarct size was approximately twice as large (80 vs. 40mm^3^, respectively) as expected from the rCBV data. Hence, brain regions with a significant reduction of rCBV as measured by functional ultrasound imaging did not fully overlap with infarct size and location, especially for the CCAo+MCAo stroke model, in contrast to what was previously observed in a thromboembolic mice model^51^.

Nevertheless, we hypothesize that functional ultrasound imaging may predict more accurately the size and location of the constituted infarct at 24hrs by merging information on the level of rCBV with analysis of hemodynamic changes associated with SDs^45,64^. Indeed, we showed that the final brain infarct was located more rostrally as compared to the initial ischemic territory measured 70mins after stroke in both models (Figure 3(c) and (d)). Additionally, we demonstrate that the SDs propagate from one point of origin in a rostrocaudal direction with a median velocity of ~6mm/min, which agrees with the literature^79–83^. Interestingly, all SDs originated within brain regions that are part of the infarct but are not ischemic in the first 70mins. It is important to note that SDs may occur up to 6hrs after stroke onset, which has been shown to cause an expansion of core-infarcted tissue^66,84^. We did not observe differences in the number of the SDs between the MCAo and CCAo+MCAo stroke during the 70mins imaging period confirming previous results, which demonstrated that the number of SDs was not directly associated with the expansion of the ischemic insult^81^. In short, our work shows the relevance of functional ultrasound imaging to better understand early hemodynamic changes in preclinical stroke research. It enlightens their high complexity, which may make stroke a heterogeneous disorder. Moreover, our results suggest that the primary somatosensory cortex in rats is highly sensitive during a stroke and becomes a hot spot for generating spreading depolarization, which may contribute to delayed infarction of this region.

We used two rat stroke models to evaluate the relevance of functional ultrasound imaging in two ischemic conditions that may recapitulate the variability of infarct size and location observed in the clinic. However, functional ultrasound imaging requires a craniotomy, especially for imaging deep brain regions, since the skull is strongly attenuating high-frequency ultrasound signals^85^. Importantly, it has been demonstrated that both neurological behavior and infarction size were significantly better in rats treated very early by decompressive craniectomy (4hrs) after endovascular MCAo^86^. Therefore, it is likely that our results may underestimate those in close skull since acute ischemia in the MCA territory may lead to cerebral edema with raised intracranial pressure and reduce collateral blood flow, which is associated with larger infarct size^87^.

Two other limitations of our study relate to i) the lack of monitoring of physiological parameters (i.e., blood pressure, blood gas analysis,...) and ii) the anesthesia. It has been demonstrated that several anesthetics have dose-specific effects on cerebral blood flow^88–91^ but also affects neurovascular coupling, autoregulation, ischemic depolarizations^91–94^, excitotoxicity, inflammation, neural networks, and numerous molecular pathways relevant for stroke outcome^91,95^. Additionally, we did not analyze the behavioral outcome, which the STAIR committee recommends for improving the translation value of preclinical stroke research^70^. These limitations could be further addressed using volumetric functional ultrasound imaging (vfUSI) that has been recently developed and based on a 2D-array transducer to acquire 3D images of brain activity in awake conditions^55^.

The versatility, ease-of-use, contrast-free, low cost, portability, and recent success in the clinic^60,96,97^ makes functional ultrasound imaging a promising tool to monitor and map large-scale and in-depth hemodynamics for a better diagnosis and prognosis of various brain vascular disorders.

## Author contributions

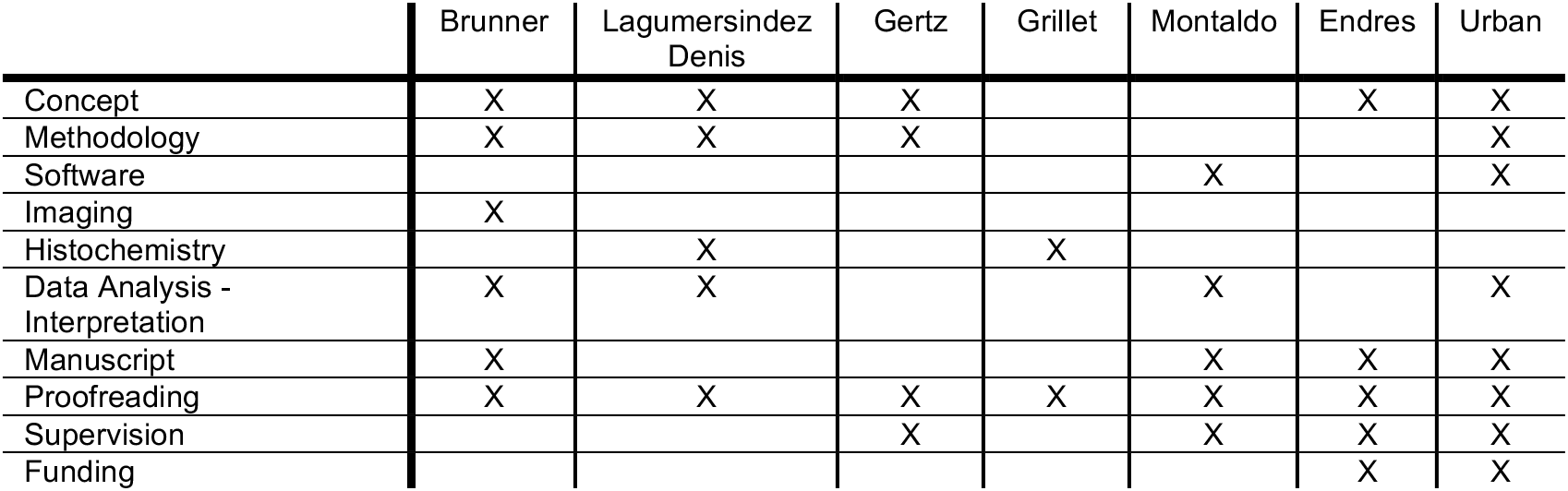

### Funding

This work is supported by grants from the Fondation Leducq (15CVD02) and KU Leuven (C14/18/099-STYMULATE-STROKE). The functional ultrasound imaging platform is supported by grants from FWO (MEDI-RESCU2-AKUL/17/049, G091719N, and 1197818N), VIB Tech-Watch (fUSI-MICE), Neuro-Electronics Research Flanders TechDev fund (3D-fUSI project). The Endres lab received funding from DFG under Germany’s Excellence Strategy – EXC-2049 – 390688087, BMBF, DZNE, DZHK, EU, Corona Foundation, and Fondation Leducq

## Acknowledgements

The authors thank all the members of the Fondation Leducq network #15CVD02 for helpful comments and discussions. We thank all NERF animal caretakers, including I. Eyckmans, F. Ooms, and S. Luijten, for their help with the management of the animals.

## Supplementary Materials

**Supplementary Figure 1.**
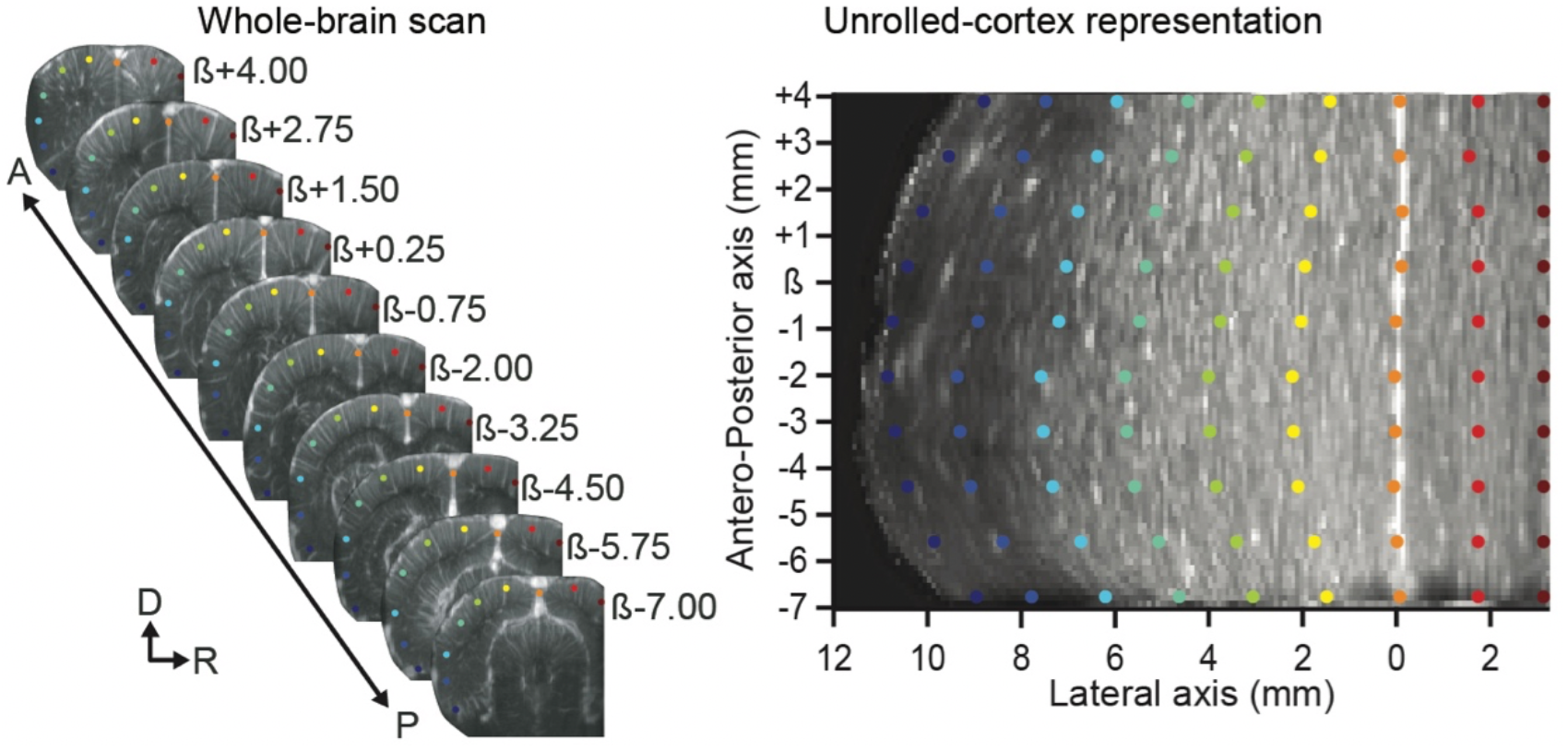
Unrolled-cortex projection.

**Supplementary Table 1.** List of the 115 brain regions extracted from a digital version of the rat Paxinos atlas organized by main anatomical structures (left) and clustered based on the average rCBV loss after the MCAo (right).

**Movie 1.** Whole-brain scans before (left) and 70mins after CCA and MCA occlusions (right).

**Movie 2.** Movie of hemodynamic changes induced by concomitant CCA and MCA occlusions observed in a single coronal plane (ß+3mm) extracted from the whole-brain scan.

**Movie 3.** Movie of hemodynamic changes induced by concomitant CCA and MCA occlusions observed with the unrolled-cortex projection.

