## Supplementary figures and images for "Brain-wide continuous functional ultrasound imaging for real-time monitoring of hemodynamics during ischemic stroke"

### SupplementaryFigure1.pdf

#### Supplementary Figure 1

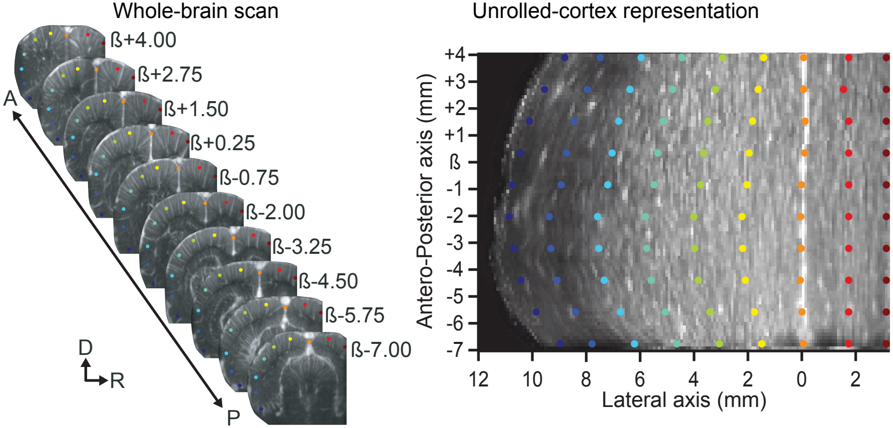
